# GWAS and genomic selection for marker-assisted development of sucrose enriched soybean cultivars

**DOI:** 10.1101/2023.04.16.537083

**Authors:** Awais Riaz, Qasim Raza, Anuj Kumar, Derek Dean, Kenani Chiwina, Theresa Makawa Phiri, Julie Thomas, Ainong Shi

**Affiliations:** Department of Crop, Soil, and Environmental Sciences, University of Arkansas, Fayetteville, AR 72701, USA; Precision Agriculture and Analytics Lab, National Centre in Big Data and Cloud Computing, Centre for Advanced Studies in Agriculture and Food Security, University of Agriculture Faisalabad, Faisalabad 38000, Pakistan; Department of Horticulture, University of Arkansas, Fayetteville, AR 72701, USA

**Keywords:** Food-grade soybean, Genomic prediction, Population genetics, Quantitative genetics, Seed sucrose concentration, Single nucleotide polymorphism (SNP), Soluble sugars

## Abstract

Sucrose concentration in soy-derived foods is becoming a seminal trait for the production of food-grade soybeans. However, limited scientific knowledge is reported on this increasingly important breeding objective. In this study, 473 genetically diverse soybean germplasm accessions and 8,477 high-quality single nucleotide polymorphisms (SNPs) markers were utilized to pinpoint genomic regions associated with seed sucrose contents through a genome-wide association study (GWAS). A total of 75 significant SNPs (LOD ≥6.0) were identified across GLM, FarmCPU and BLINK models, including four stable and novel SNPs (Gm03_45385087_ss715586641, Gm06_10919443_ss715592728, Gm09_45335932_ss715604570 and Gm14_10470463_ss715617454). Gene mining near 20 kb flanking genomic regions of four stable SNPs identified 23 candidate genes with the majority of them highly expressed in soybean seeds and pod shells. A sugar transporter encoding major facilitator superfamily gene (*Glyma*.*06G132500*) showing the highest expression in pod shells was also identified. Moreover, selection accuracy, efficiency and favorable alleles of 75 significantly associated SNPs were estimated for their utilization in soybean breeding programs. Furthermore, genomic predictions with three different scenarios revealed better feasibility of GWAS-derived SNPs for selection and improvement of seed sucrose concentration. These results could facilitate plant breeders in marker-assisted breeding and genomic selection of sucrose-enriched food-grade soybean cultivars for the global soy-food industry.

## Introduction

Soybean (*Glycine max* L.) is an important legume crop cultivated worldwide for food, feed, industrial and bioenergy purposes (Qiu et al. 2011). It is integrated as a rotational crop in cropping schemes due to nitrogen-fixing properties, which aids in more sustainable agricultural practices (Lee et al. 2015). The demand for soy-derived foods is increasing over the past few decades due to popularity of plant-based diets (Lynch et al. 2018). On a dry matter basis, soybean seeds contain approximately 40% proteins, 35% carbohydrates, 20% oils and 5% ash (Krober and Cartter 1962). Among carbohydrates, about 40% are soluble sugars which are pivotal for soybean nutritional quality and further comprised of 5% sucrose, 3% stachyose and 1.5% raffinose (Wilson 2016). Soybean seeds also contain other minor soluble sugars such as fructose, glucose and galactose (Hymowitz et al. 1972). Soluble sugars, especially sucrose, substantially affect the nutritional quality of soy-derived foods including edamame, natto, soymilk, tempeh and tofu (Taira et al. 1990; Poysa and Woodrow 2002; Hou et al. 2009). High sucrose concentration improves the sweetness and flavor of soy foods. Additionally, sucrose is involved in various metabolic and biosynthetic processes, plays a vital role in growth and development, coordinates relationship between plant sources and sinks, and responds to different abiotic stresses (Farrar et al. 2000; Ruan et al. 2010; Ruan 2012; Du et al. 2020). Thus, the paramount roles of sucrose in higher plants cannot be underestimated, especially in nutritional quality improvement of plant-based foods.

Food-grade soybean cultivars are chosen based on physical, chemical and processing quality attributes of seeds (Rao et al. 2002). To successfully meet the food-grade quality standards set by the *Ontario Soybean and Canola Committee* (OSACC), soybean seeds must contain 6.0–8.0% sucrose content (OSACC 2020). The OSACC classifies soybean cultivars based on seed sucrose contents as low (<6.4%), moderate (6.4–7.0%) and high (>7.0%). Seed sucrose concentration varies significantly among cultivars and is highly influenced by environment and agronomic management practices (Bellaloui et al. 2011; Li et al. 2012). Therefore, breeding efforts received little attention for sucrose improvement as compared with other agronomic and quality traits, as later are comparatively more valuable to commodity-grade soybeans. Nevertheless, with shifts in demand for plant-based foods, genetic breeding for sucrose-enriched seeds is becoming a major objective of soybean breeders.

Linkage mapping has been effectively utilized to identify quantitative trait loci (QTL) associated with soybean seed sucrose contents. Maughan et al. (2000) identified 17 QTLs related to sucrose contents using a recombinant inbred line (RIL) population. Kim et al. (2005) & (2006) reported six sucrose-contributing QTLs on chromosomes 2, 11, 12, 16, and 19 using a RIL population derived from a cross between Keunolkong and Shinpaldalkong soybean cultivars. Skoneczka et al. (2009) reported a major QTL on chromosome 6 in an interval of Sat_213 and Satt_643 molecular markers using two F_2_ segregation populations, which could explain 76% of the total genetic variation in sucrose concentration. Maroof and Buss (2011) detected a large effect QTL linked with sucrose and stachyose contents on chromosome 11. Wang et al. (2014) reported one QTL for sucrose and raffinose contents in a set of 170 F_2:3_ derived RILs. Zeng et al. (2014) identified three sucrose-contributing QTLs from an F_2_ mapping population derived from contrasting parents MFS-553 (low sucrose) and PI 243545 (high sucrose). These QTLs were mapped on chromosomes 5, 9, and 16 and could explain 46%, 10% and 8% variations in the total sugar content, respectively. Akond et al. (2015) detected 14 QTLs significantly associated with soluble sugar contents including three sucrose-related QTLs in a set of 92 F_5:7_ derived population. Recently, Patil et al. (2018) identified four QTLs on chromosomes 6, 8, 16, and 20 using a RIL population derived from *G. max* (Williams 82) and *G. soja* (PI 483460B) and reported a major QTL for sucrose contents (*qSuc_08*) on chromosome 8. However, all these family-based QTL mappings relied upon biparental populations which offer limited resolution and precision, requiring the use of high-throughput and accurate modern approaches.

As an alternative to linkage analysis, GWAS has been used to elucidate the molecular basis of complex traits. Advances in genome sequencing techniques have accelerated the resolution and accuracy of GWAS in soybean (Li et al. 2015). Next-generation sequencing enables the GWAS to precisely uncover the genetic basis of quantitative traits and their underlying putative QTLs/SNPs as compared with traditional QTL mapping techniques (Korte and Farlow 2013; Fang et al. 2017). The application of low-cost and high throughput sequencing approaches such as genotyping-by-sequencing (GBS) with advancement in computational analysis has enabled to design molecular markers that could help to improve complex traits with high selection accuracy and efficiency. The use of diverse genotypic panels in GWAS provides comprehensive information on allelic diversity and single nucleotide polymorphisms (SNPs) in different genetic backgrounds that may not be possible with a biparental population (Heffner et al. 2009). Using GWAS, several studies on the improvement of soybean agronomic and quality traits have been reported (Hwang et al. 2014; Zhou et al. 2015; Cao et al. 2017; Khan et al. 2019; Yang et al. 2020; Lee et al. 2021). However, to date, only a few studies involving the GWAS approach have been conducted to identify genomic regions associated with sucrose content in soybean seeds (Sui et al. 2020; Ficht et al. 2022; Xu et al. 2022; Lu et al. 2022).

In this study, a GWAS was performed using 473 diverse soybean germplasm accessions and 8,477 high-quality SNPs to pinpoint stable and large-effect genomic regions associated with seed sucrose contents. Selection accuracy, efficiency and favorable alleles of significant SNPs, as well as candidate genes were also identified for marker-assisted breeding and genomic selection of sucrose enriched soybean cultivars. The improved knowledge would facilitate towards development of food-grade cultivars and enhance the global soy-derived food market.

## Materials and methods

### Plant materials

The United States Department of Agriculture (USDA) soybean germplasm panel comprising 5,483 accessions along with seed sucrose concentration data were retrieved from the U.S. National Plant Germplasm System (https://npgsweb.ars-grin.gov/gringlobal/methodaccession?id1=51073&id2=494186, accessed on January 10, 2023). The USDA germplasm panel consisted of breeding material, elite lines, approved cultivars, obsolete cultivars and landraces. Among this panel, 473 accessions with 100 seed weights of >23 grams were filtered out and chosen for GWAS (**Table S1**). The selected accessions mainly originated from 11 countries including China, France, Japan, Nepal, North Korea, Russian Federation, South Korea, Sweden, Taiwan and the United States. The USDA–ARS germplasm curation staff and their collaborators conducted field experiments at various location across several years to record the seed weight of soyabean germplasm panel. Sucrose concentration in this germplasm panel was determined by following standard sucrose extraction and quantification methods (Choung 2010; Teixeira et al. 2012) and detail are provided in GRIN (https://npgsweb.ars-grin.gov/gringlobal/method?id=494186)

### Genotyping

Genomic DNA was isolated from fresh leaves of selected germplasm panel using DNeasy 96 Plant Kit (QIAGEN, Valencia, CA), followed by genotyping with Illumina Golden Gate SNP assay. The panel was genotyped with Soy5K SNP Infinium Chips (Song et al., 2013) consisting of 42,291 SNPs distributed across 20 soybean chromosomes were downloaded from https://www.soybase.org/snps/download.php. The density and distribution map of SNPs on each soybean chromosome was drawn using a CMplot package (Yin et al. 2021) and shown in **Fig. S1**. SNPs with >10% missing & heterozygosity data, <5% minor allele frequency and tri-allelic status were removed, keeping only 8,477 high-quality SNPs for further analyses.

### Population genetics analyses

Principle component analysis (PCA) and neighbor-joining phylogenetic tree analysis were conducted through the Genomic Association and Prediction Integrated Tool version 3 (GAPIT3) (Wang and Zhang 2021) implemented in R (v4.1.3; R Core Team 2022) with default settings. The possible population structure was inferred based on block principal pivoting method using the LEA package (Frichot and François 2015) in R. The simulations were performed using genotypic information of 8,477 high-quality SNPs by setting 1,000 repetitions, 100 alpha, 100 iterations and number of populations (K) from 2 to 10. The cross-entropy inference was set at a threshold of 5% and a Q-matrix threshold of >0.5 for subsequent population structure analysis of 473 USDA soybean germplasm accessions. Moreover, a similar attempt was conducted with the whole set of SNPs to find possible discordance in population structure derived from high-quality SNPs, however, a non-significant difference in population structure was observed (data not shown). Hence, only high-quality SNP set-derived population structure was considered for this study.

### Association analysis

GWAS was performed to find SNPs significantly associated with seed sucrose concentration using generalized linear model (GLM) (Nelder and Wedderburn 1972), mixed linear model (MLM) (Zhang et al. 2010), fixed and random model circulating probability unification (FarmCPU) (Liu et al. 2016), and Bayesian-information and linkage-disequilibrium iteratively nested keyway (BLINK) (Huang et al. 2019) models implemented in GAPIT3 (Wang and Zhang 2021). In this study, a comparatively strict logarithmic of odds (LOD ≥6.0) threshold based upon the Bonferroni correction test (α = 0.05) (Bland and Altman 1995) was set to determine highly significant associations.

### Candidate gene analysis

Candidate genes were mined within 20 kilobase pairs (kbp) flanking genomic regions on either side of four stable and significant SNPs identified from GWAS. The soybean reference genome assembly (Gmax_Wm82_a2_v1) (Schmutz et al. 2010) was searched for retrieval of annotated and protein-encoding candidate genes. Functional annotation and *in*-*silico* gene expressions were retrieved from SoyBase (https://soybase.org) (Grant et al. 2010) and Soybean eFP Browser (http://bar.utoronto.ca/efpsoybean/cgi-bin/efpWeb.cgi).

### Favorable SNP alleles, selection accuracy and selection efficiency

Favorable alleles of GWAS-derived 75 significant SNPs were determined from genotypic and phenotypic data of 114 contrasting soybean accessions containing high and low (57 each) seed sucrose contents. Similarly, SNP selection accuracy and efficiency were computed by following previously reported formulas (Shi et al. 2016; Ravelombola et al. 2019).

Selection accuracy = 100 x [(Number of genotypes having high sucrose with favorable SNP allele) / (Number of genotypes having high sucrose with favorable SNP allele + Number of genotypes having low sucrose with favorable SNP allele)]

Selection efficiency = 100 x [Number of genotypes having high sucrose with favorable SNP allele / Total number of genotypes having favorable SNP allele]

### Genomic selection/prediction

Genomic selection/prediction was computed based on genomic estimated breeding values (GEBVs) of 473 selected germplasm panel. GEBVs were estimated using nine genomic prediction models (BA, BB, BL, BRR, cBLUP, gBLUP, RF, rrBLUP and SVM) with five different training to testing set ratios (50/50, 67/33, 75/25, 83/17 and 90/10) and whole-set versus GWAS-derived SNPs. For GP estimation using different models, ridge regression best linear unbiased prediction (rrBLUP) package (Endelman 2011) was implemented in R, genomic BLUP (gBLUP) and compressed BLUP (cBLUP) implemented in GAPIT3 (Wang and Zhang 2021), Bayes A (BA), Bayes B (BB), Bayes LASSO (BL) and Bayes ridge regression (BRR) implemented in BGLR (Pérez and De Los Campos 2014), random forest (RF) implemented in randomForest R package (Breiman 2001) and support vector machine (SVM) implemented in kernlab packages (Karatzoglou et al. 2023). Prediction accuracy of tested GP models was computed using the average Pearson’s correlation coefficient (r) between GEBV estimates from training and testing sets (Zhang et al. 2016). The averaged r-values were estimated from 100 repetitions of five above-mentioned training to testing sets and boxplots were generated using the ggplot2 package (Wickham 2016).

## Results

### Genetic diversity among USDA soybean germplasm

The USDA soybean germplasm consisted of a hefty collection of diverse accessions collected from all over the world. Among this collection, 473 accessions with 100 seed weights >23 g were chosen for this study (**Table S1**). The majority of the accessions originated from South Korea, Japan and China with marginal contributions from eight other countries. The chosen accessions included advanced breeding material, elite lines, cultivars, landraces and wild relatives. Sucrose in germplasm collection varied from 0.1% to 10.5% with a median value of 4.3% (**Fig. 1A**). Frequency distribution plot indicated a near symmetrical distribution of sucrose contents and germplasm accessions. Wider genetic diversity in Japanese, South Korean, Chinees and Russian accessions was observed as sucrose in these countries accessions ranged from 0.4–10.5%, 1.2–10.2%, 0.1–6.5% and 1.2–6.8% respectively (**Fig. 1B**), indicating Asia as edamame soybean centre of origin and further improvement. Two accessions (PI398768 and PI229343) with >10% sucrose contents were recognized from this diverse germplasm set (**Table S1**), which could be exploited in soybean breeding programs as donor parents for the development of higher-yielding and more nutritious cultivars. Hence, the germplasm collection was genetically diverse for conducting genome-wide association and genomic selection studies.

**Fig. 1.**
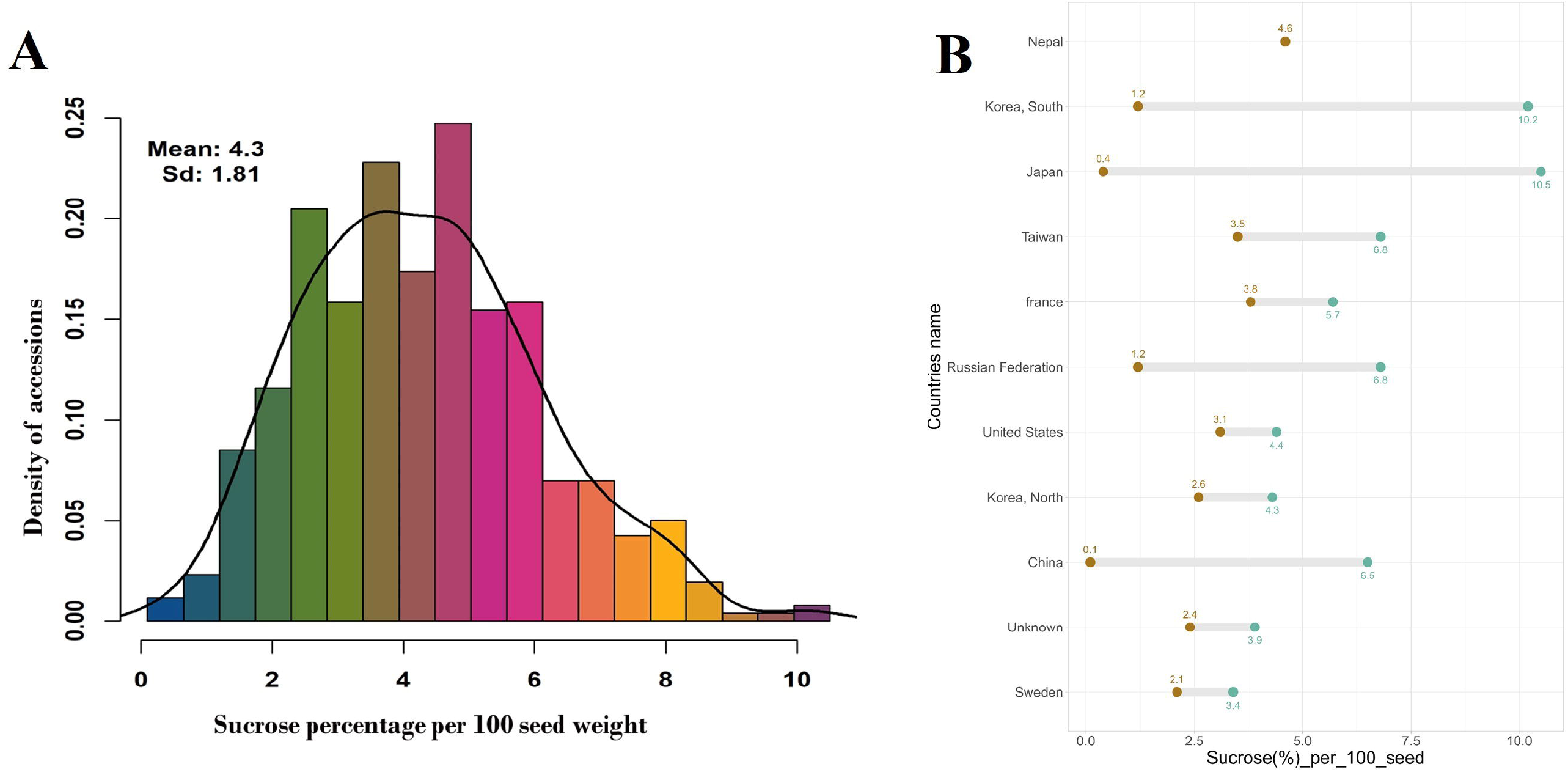
Genetic diversity for sucrose contents among USDA soybean germplasm. (**A**) Frequency distribution and (**B**) range of sucrose contents in soybean seeds among 473 accessions originating from different countries.

### Population structure, PCA and phylogenetic analyses

By subjecting the genotypic information of 8,477 high-quality SNPs set, population structure, PCA and phylogenetic analyses were carried out for 473 soybean accessions selected from the USDA germplasm. An optimal number of three ancestral sub-papulations or clusters were inferred with K value 2–10 and Q-matrix threshold >0.5 using LEA package in R (**Table S2**). The projected population structure is shown in **Fig. 2A**. A similar attempt utilizing the whole 42,291 SNPs set yielded a non-significant difference in structure results when compared with high-quality SNPs set-derived population structure (data not shown). Cluster I, II and III consisted of 41 (8.67%), 167 (35.31%) and 238 (50.32%) accessions with major contributions from China, South Korea, and Japan, respectively (**Fig. 2A** and **Table S2**).

**Fig. 2.**
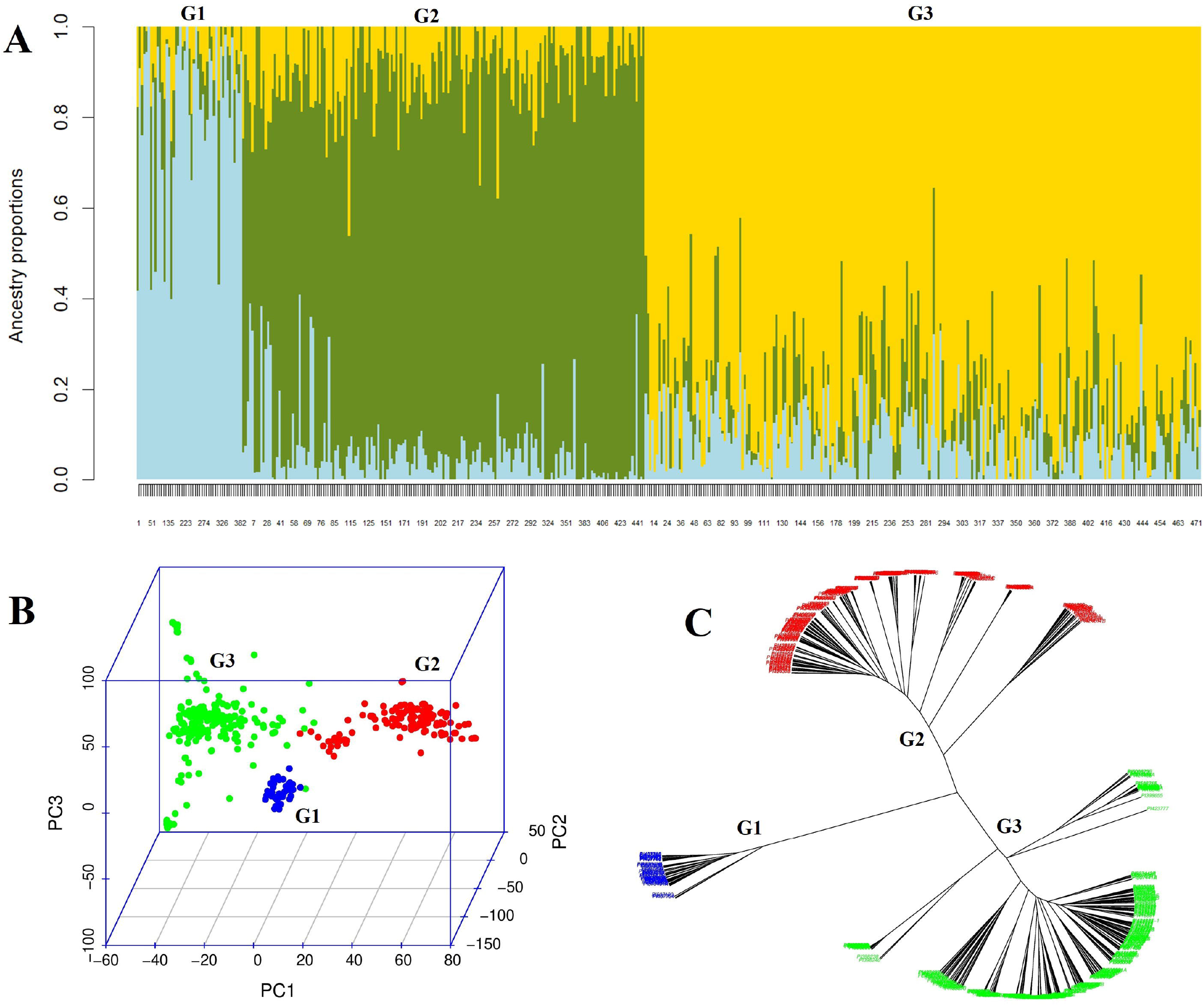
Population diversity analysis of 473 USDA soybean germplasm accessions. (**A**) Classification of soybean accessions into three sub-populations using LEA-package in R. Numbers on x- and y-axes indicate different accessions and ancestry proportions, respectively. Sub-populations are indicated with G1, G2 and G3. (**B**) Principal component analysis of soybean accessions. Different axes represent three principal components (PC1, PC2 and PC3) and variances explained by these PCAs. Sub-populations/clusters are indicated with G1, G2 and G3. (**C**) Neighbor-joining phylogenetic tree of 473 soybean accessions with three distinct sub-populations/clusters/groups.

Similarly, PCA and phylogenetic analyses clearly distributed 473 soybean accessions into three distinct clusters or groups by using the high-quality SNPs set information subjected to GAPIT 3 (**Figures 2B** and **2C**). Collectively, these results support genetic diversity results and indicate that sub-populations or clusters have stable boundaries with well-defined genetic backgrounds.

### GWAS for sucrose contents

GWAS was conducted to pinpoint high-quality SNPs significantly associated with sucrose contents in USDA soybean germplasm. GLM, MLM, FarmCPU and BLINK models were tested by GAPIT 3. Combined and individual Manhattan plots identified a total of 75 SNPs significantly associated (LOD ≥6.0) with sucrose contents in seeds (**Figs. 3A** and **S2A–S5A**). The number of significant SNPs varied from 6 to 63 among GLM, FarmCPU and BLINK, without single SNP contribution by MLM, and unevenly distributed on 13 out of 20 soybean chromosomes (**Table S3**). The observed LOD distributions [−log10(p)] in QQ-plots also showed deviations from expected LOD distributions (**Figs. 3B** and **S2B–S5B**) confirming a significant association of identified SNPs with sucrose contents in USDA germplasm. Moreover, this study also identified a single overlapping SNP (Gm03_45385087_ss715586641) with LOD >6.0 across GLM, FarmCPU and BLINK models (**Fig. 3A**), indicating the presence of a highly stable large effect quantitative trait loci (QTL) significantly associated with sucrose contents on chromosome 3. Furthermore, three other overlapping significant SNPs were also pinpointed by GLM and FarmCPU (Gm06_10919443_ss715592728 and Gm09_45335932_ss715604570) or GLM and BLINK (Gm14_10470463_ss715617454) models, revealing preponderance of stable and large effect QTLs for sucrose contents on chromosomes 6, 9 and 14 (**Table 1**).

**Fig. 3.**
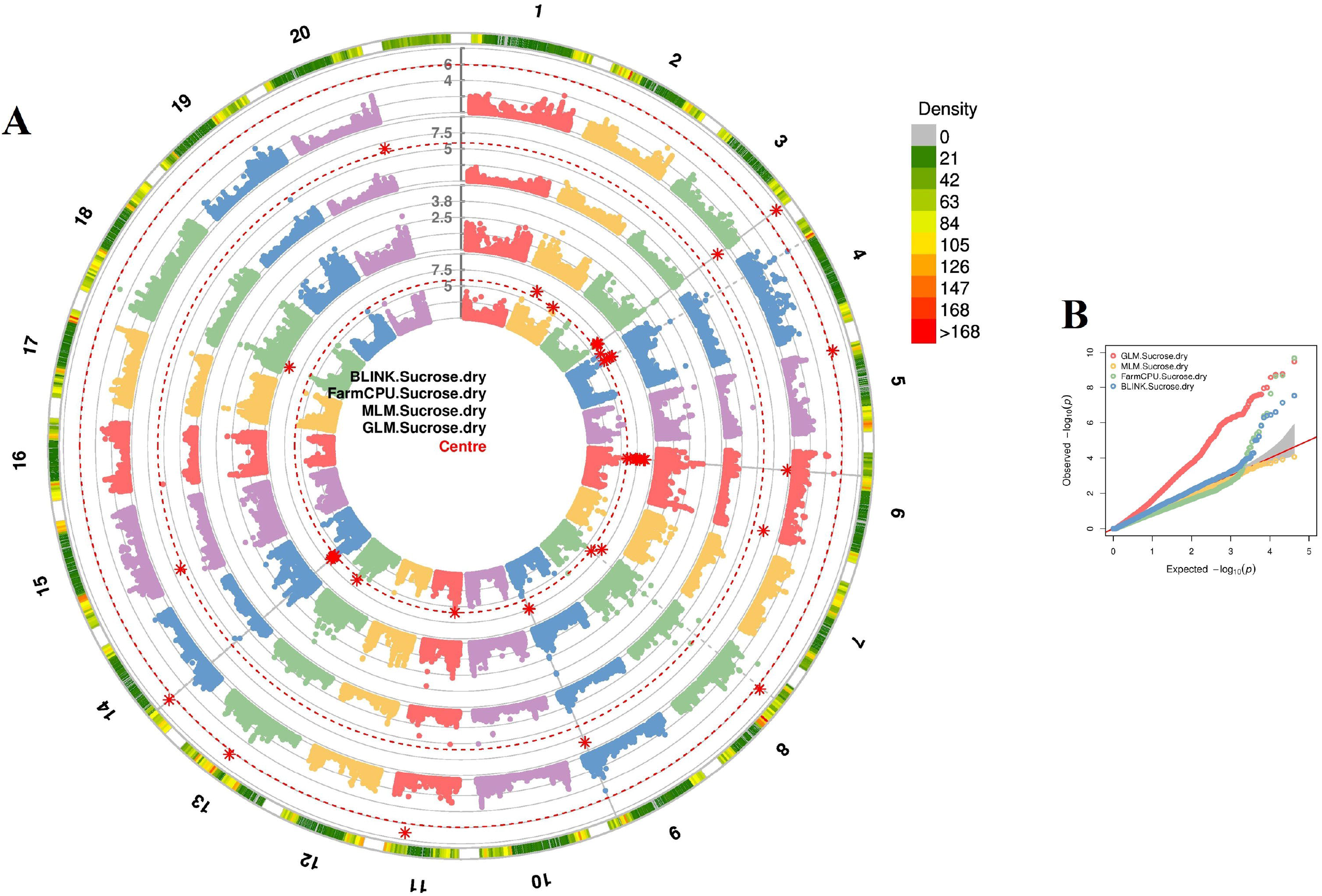
Genome-wide association study for high sucrose contents in soybean seeds. (**A**) A combined Manhattan plot with GLM, MLM, FarmCPU and BLINK models. The outermost band indicates 20 soybean chromosomes and a density of high-quality SNPs on these chromosomes. Four inner bands represent the distribution of SNPs on respective soybean chromosomes using BLINK, FarmCPU, MLM and GLM models. Significant LOD threshold (≥6.0) and SNPs across different models are indicated with dotted red circles and asterisk marks, respectively. (**B**) A combined QQ-plot showing observed vs expected LOD (−log(P-value)) values across four tested models.

**Table 1.**
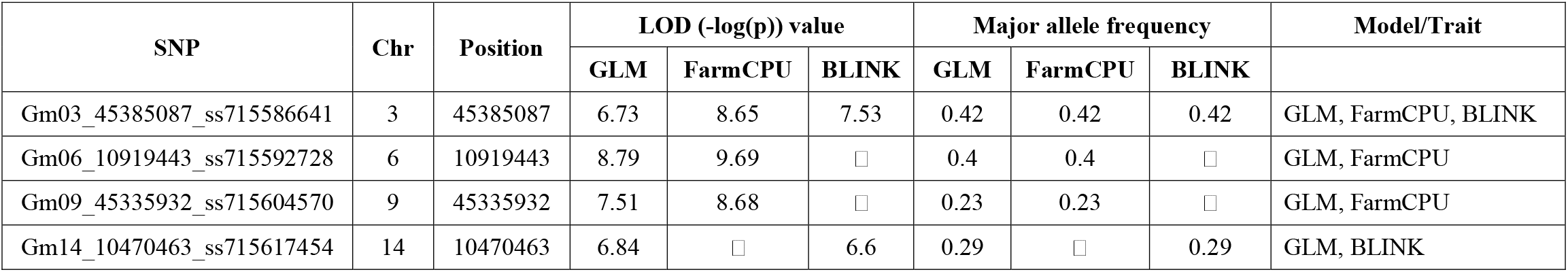
Significant and overlapping SNPs associated with sucrose contents in USDA soybean germplasm

GLM identified 63 SNPs on chromosomes 2, 3, 4, 6, 8, 9, 11, 13, 14 and 18 (**Fig. S2A**) with LOD and MAF ranging from 6.0–9.49 and 0.084–0.490, respectively (**Table S3**). Whereas MLM was unable to find even a single SNP with LOD ≥6.0, however, few SNPs were associated with sucrose contents at a lower LOD value (≥4.0) (**Fig. S3A**). FarmCPU and BLINK identified six SNPs each on chromosomes 3, 5, 6, 8, 9, 11, 13, 14 and 15 (**Figs. S4A** and **S5A**). The LOD and MAF of FarmCPU-derived SNPs ranged from 6.38–9.69 and 0.064–0.425, respectively. Likewise, BLINK-derived SNPs had LOD and MAF in the range of 6.28–7.53 and 0.022–0.425, respectively (**Table S3**). In this study, identification of at least four stable and large-effect SNPs/QTLs significantly associated with sucrose contents in seeds (**Table 1**) could provide a basis for marker-assisted development of high sucrose-containing soybean genotypes.

### Candidate genes associated with sucrose contents

Candidate genes for sucrose contents were mined from 20 kb flanking genomic regions of four stable and large-effect SNPs through the Phytozome database. A total of 23 protein-encoding candidate genes were identified whose start or end positions lay within 20 kb genomic regions (**Table 2**). Among these, 9 genes code for hypothetical proteins with unknown functions, whereas 3 genes code for P-loop containing nucleoside triphosphate hydrolases. Identified genes from chromosome 3 code for cytochrome B5 isoform E, auxin-responsive protein, Rho GTPase activating protein with PAK-box and DHHC-type zinc finger family proteins with moderate to highest expressions in soybean seeds and pod shells, along with other vegetative tissues. Similarly, candidate genes positioned on chromosome 6 code for sugar transporter/major facilitator superfamily protein, rhodanese/cell cycle control phosphatase superfamily protein and P-loop containing nucleoside triphosphate hydrolase superfamily proteins with moderate to highest expressions in seeds and pod shells. Whereas candidate genes from chromosome 9 code for purple acid phosphatase 10, chromatin remodeling 31, NAD(P)-linked oxidoreductase superfamily protein, prefoldin 6 and RNI-like superfamily proteins with only three showing moderate to highest expressions in reproductive, as well as, in vegetative tissues. However, both of the genes positioned on chromosome 14 code for hypothetical proteins and only one is highly expressed in root tissues. Putative functions of all these candidate genes included energy production for conformational changes, transmembrane transport of materials and regulation of proteins localization, stability and activity. Interestingly, a sugar transporter encoding major facilitator superfamily gene (*Glyma*.*06G132500*) was also identified in this study which showed highest expressions in soybean pod shells at 10– 17 days after fertilization (DAF) (Severin et al. 2010). None of the four stable SNPs was positioned within the genic region of any candidate gene, as all four SNPs were positioned at <5–25 kb distance from start or end position of all candidate genes (**Table 2**). Further detailed studies are required to explore the putative roles of identified candidate genes in sucrose biosynthesis, transport, and storage.

**Table 2.**
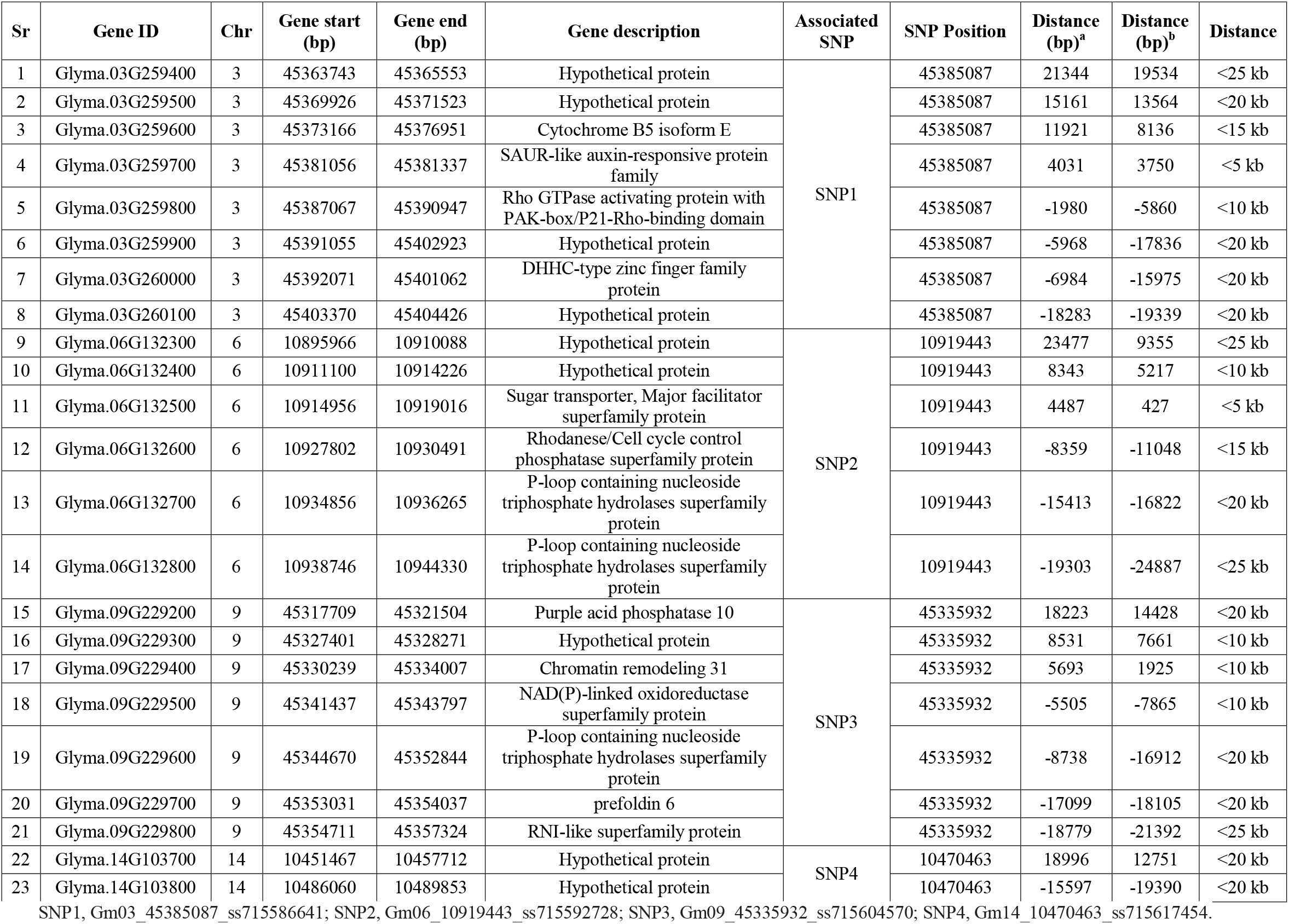
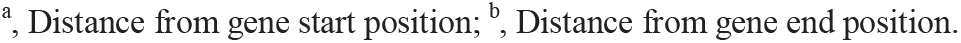
Candidate genes associated with sucrose contents in soybean seeds

| Sr | Gene ID | Chr | Gene start (bp) | Gene end (bp) | Gene description | Associated SNP | SNP Position | Distance (bp)<sup>a</sup> | Distance (bp)<sup>b</sup> | Distance |
| --- | --- | --- | --- | --- | --- | --- | --- | --- | --- | --- |
| 1 | Glyma.03G259400 | 3 | 45363743 | 45365553 | Hypothetical protein | SNP1 | 45385087 | 21344 | 19534 | <25 kb |
| 2 | Glyma.03G259500 | 3 | 45369926 | 45371523 | Hypothetical protein |  | 45385087 | 15161 | 13564 | <20 kb |
| 3 | Glyma.03G259600 | 3 | 45373166 | 45376951 | Cytochrome B5 isoform E |  | 45385087 | 11921 | 8136 | <15 kb |
| 4 | Glyma.03G259700 | 3 | 45381056 | 45381337 | SAUR-like auxin-responsive protein family |  | 45385087 | 4031 | 3750 | <5 kb |
| 5 | Glyma.03G259800 | 3 | 45387067 | 45390947 | Rho GTPase activating protein with PAK-box/P21-Rho-binding domain |  | 45385087 | -1980 | -5860 | <10 kb |
| 6 | Glyma.03G259900 | 3 | 45391055 | 45402923 | Hypothetical protein |  | 45385087 | -5968 | -17836 | <20 kb |
| 7 | Glyma.03G260000 | 3 | 45392071 | 45401062 | DHHC-type zinc finger family protein |  | 45385087 | -6984 | -15975 | <20 kb |
| 8 | Glyma.03G260100 | 3 | 45403370 | 45404426 | Hypothetical protein |  | 45385087 | -18283 | -19339 | <20 kb |
| 9 | Glyma.06G132300 | 6 | 10895966 | 10910088 | Hypothetical protein | SNP2 | 10919443 | 23477 | 9355 | <25 kb |
| 10 | Glyma.06G132400 | 6 | 10911100 | 10914226 | Hypothetical protein |  | 10919443 | 8343 | 5217 | <10 kb |
| 11 | Glyma.06G132500 | 6 | 10914956 | 10919016 | Sugar transporter, Major facilitator superfamily protein |  | 10919443 | 4487 | 427 | <5 kb |
| 12 | Glyma.06G132600 | 6 | 10927802 | 10930491 | Rhodanese/Cell cycle control phosphatase superfamily protein |  | 10919443 | -8359 | -11048 | <15 kb |
| 13 | Glyma.06G132700 | 6 | 10934856 | 10936265 | P-loop containing nucleoside triphosphate hydrolases superfamily protein |  | 10919443 | -15413 | -16822 | <20 kb |
| 14 | Glyma.06G132800 | 6 | 10938746 | 10944330 | P-loop containing nucleoside triphosphate hydrolases superfamily protein |  | 10919443 | -19303 | -24887 | <25 kb |
| 15 | Glyma.09G229200 | 9 | 45317709 | 45321504 | Purple acid phosphatase 10 | SNP3 | 45335932 | 18223 | 14428 | <20 kb |
| 16 | Glyma.09G229300 | 9 | 45327401 | 45328271 | Hypothetical protein |  | 45335932 | 8531 | 7661 | <10 kb |
| 17 | Glyma.09G229400 | 9 | 45330239 | 45334007 | Chromatin remodeling 31 |  | 45335932 | 5693 | 1925 | <10 kb |
| 18 | Glyma.09G229500 | 9 | 45341437 | 45343797 | NAD(P)-linked oxidoreductase superfamily protein |  | 45335932 | -5505 | -7865 | <10 kb |
| 19 | Glyma.09G229600 | 9 | 45344670 | 45352844 | P-loop containing nucleoside triphosphate hydrolases superfamily protein |  | 45335932 | -8738 | -16912 | <20 kb |
| 20 | Glyma.09G229700 | 9 | 45353031 | 45354037 | prefoldin 6 |  | 45335932 | -17099 | -18105 | <20 kb |
| 21 | Glyma.09G229800 | 9 | 45354711 | 45357324 | RNI-like superfamily protein |  | 45335932 | -18779 | -21392 | <25 kb |
| 22 | Glyma.14G103700 | 14 | 10451467 | 10457712 | Hypothetical protein | SNP4 | 10470463 | 18996 | 12751 | <20 kb |
| 23 | Glyma.14G103800 | 14 | 10486060 | 10489853 | Hypothetical protein |  | 10470463 | -15597 | -19390 | <20 kb |

SNP1, Gm03\_45385087\_ss715586641; SNP2, Gm06\_10919443\_ss715592728; SNP3, Gm09\_45335932\_ss715604570; SNP4, Gm14\_10470463\_ss715617454.
<sup>a</sup>, Distance from gene start position; <sup>b</sup>, Distance from gene end position.

### Marker-assisted selection accuracy and efficiency of GWAS-derived SNPs

In this study, GWAS-derived SNPs selection accuracy and efficiency along with favorable alleles were also estimated for marker-assisted selection of high sucrose-containing soybean genotypes. Large variations in SNP selection accuracy and efficiency were observed, as these indices varied from 53%–100% and 3%–50% with averages of 70% and 39%, respectively (**Table S4**). At least five significant SNPs (Gm04_9297161_ss715589625¬_C_T, Gm04_9412759_ss715589652_A_A, Gm04_9510512_ss715589670_C_T,

Gm04_9581724_ss715589687_C_T and Gm05_1375337_ss715592485_A_G) exhibited 100% accuracy but inadequate efficiency (3%–17%). As expected, high selection accuracy (>66%) and efficiency (>40%) were observed for all four overlapping and significant SNPs, except for Gm09_45335932_ss715604570_C_T which showed moderate efficiency (24.5%). The favorable alleles for high sucrose contents of four overlapping SNPs including Gm03_45385087_ss715586641_A_C, Gm06_10919443_ss715592728_C_T, Gm09_45335932_ss715604570_C_T and Gm14_10470463_ss715617454_G_T were found to be A, T, C and T, respectively (**Table S4**). Overall, these results indicate that GWAS-derived SNPs offer moderate to high selection accuracy but relatively lower selection efficiency, however, at least three overlapping and stable SNPs offer comparatively higher accuracy and efficiency. Favorable alleles of these stable and significant SNPs could be exploited through marker-assisted selection for screening and development of high sucrose-containing soybean genotypes.

### Genomic prediction of sucrose contents

#### Genomic prediction using different training to testing set ratios and GP models

GP was performed by subjecting five different ratios of training/testing sets (50/50, 67/33, 75/25, 83/17 and 90/10), nine GP models (BA, BB, BL, BRR, cBLUP, gBLUP, RF, rrBLUP and SVM) and the whole SNPs set, making a total of 45 combinations. The averaged Pearson correlation coefficients (r□100) along with standard errors of 100 individual runs between GEBVs and sucrose contents for 45 combinations are displayed in **Fig. 4**. Five training/testing sets had similar but not identical averaged r values across all nine GP models, except cBLUP with slightly lower r values. Increasing the training/testing ratio marginally improved selection accuracy across all tested GP models, however, the SE was also enlarged. The averaged r value across all tested models was 0.49 and ranged from 0.34–0.52 (**Table S5**). Likewise, 90/10 ratio set had the highest averaged r value (0.51) and varied from 0.36– 0.55. In general, four GP models (BB, BRR, gBLUP and rrBLUP) had the highest selection accuracy values over all other models with cBLUP having the lowest accuracy values (**Fig. 4**). Overall, these results indicate that genomic selection in soybean for sucrose contents can be carried out using any training/testing set and GP model (except for cBLUP).

**Fig. 4.**
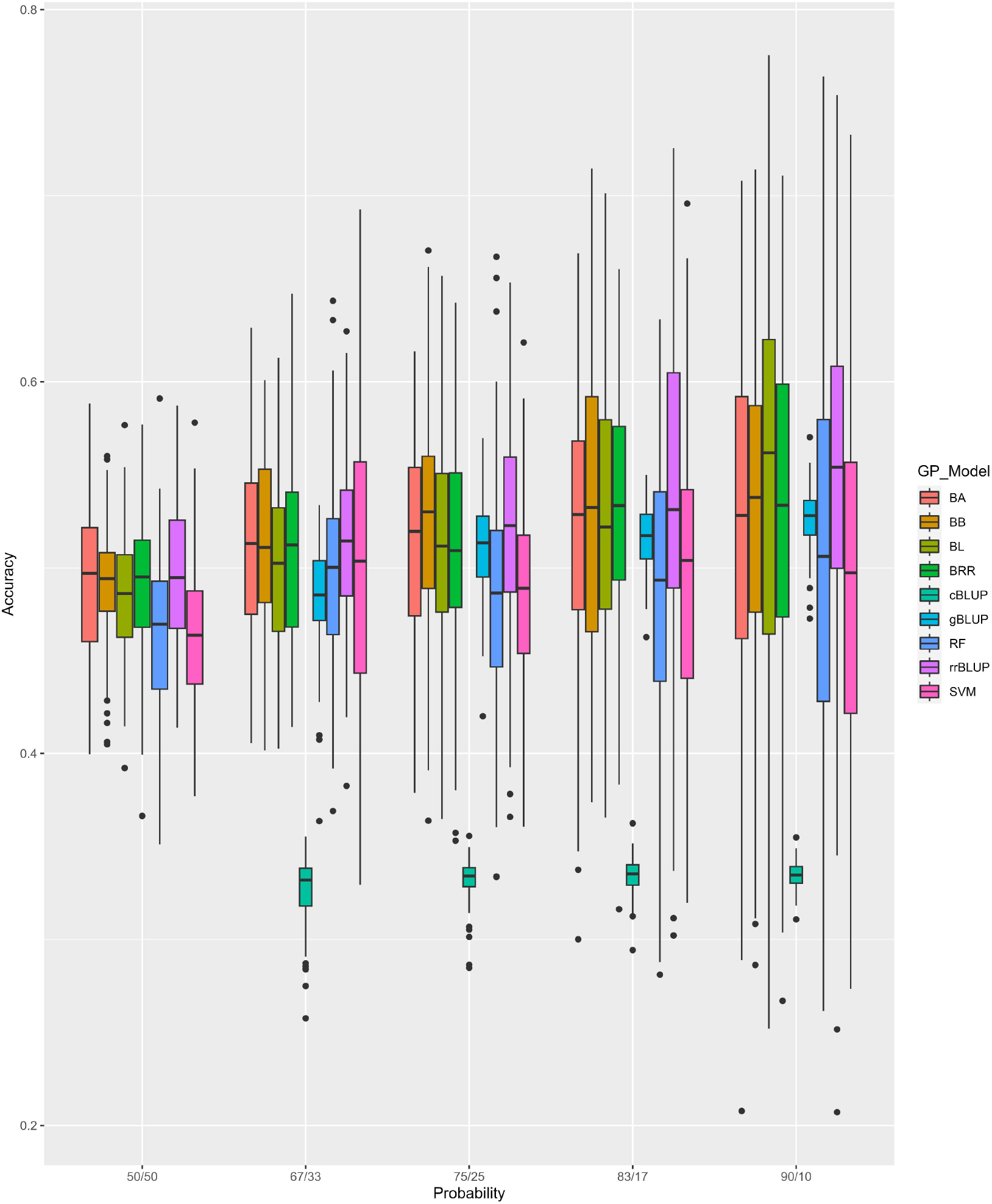
Genomic prediction of high sucrose contents using different training/testing set ratios and GP models.

#### Genomic prediction using whole set and GWAS-derived SNPs

GP was also completed between the whole set and GWAS set SNPs to explore selection efficiency of identified SNPs significantly associated with sucrose contents. Significantly higher prediction accuracy values were observed for GWAS set SNPs across all GP models as compared with whole set SNPs (**Fig. 5**). The r-values of GWAS-derived SNPs ranged from 0.44 to 0.63 with a mean value of 0.60, which are 16% to 33% higher than r-values of whole set SNPs. On average, prediction accuracy was improved by 22% with GWAS-set SNPs (**Table 3**). Among GP models, the highest increase in prediction accuracy using GWAS set SNPs was observed with cBLUP (33%), followed by BL (25%) and gBLUP (24%), whereas the lowest increase was found in rrBLUP (16%) and RF (17%). However, despite the highest increase in prediction accuracy with cBLUP, its r-value was significantly lower than r-values of all other GP models (**Fig. 5**), indicating that this model is less efficient for genomic selection of sucrose contents as compared with all other models. Collectively, these results highlight that the prediction accuracy of GWAS-derived SNPs set is high and these SNP markers could be utilized for genomic selection/prediction of high sucrose contents in soybean seeds.

**Table 3.** Genomic prediction for high sucrose contents using whole set and GWAS-derived SNPs

| GP Model | Whole_set_SNPs | GWAS_set_SNPs | Average | Change in accuracy |
| --- | --- | --- | --- | --- |
| | $r_{100}$ | $r_{100}$ | (GP model) | |
| BA | 0.512 | 0.630 | 0.571 | 23% |
| BB | 0.515 | 0.626 | 0.571 | 22% |
| BL | 0.504 | 0.63 | 0.567 | 25% |
| BRR | 0.507 | 0.626 | 0.567 | 23% |
| SVM | 0.499 | 0.607 | 0.553 | 22% |
| RF | 0.498 | 0.58 | 0.539 | 17% |
| rrBLUP | 0.514 | 0.595 | 0.554 | 16% |
| gBLUP | 0.510 | 0.633 | 0.572 | 24% |
| cBLUP | 0.332 | 0.440 | 0.386 | 33% |
| Average | 0.490 | 0.600 | 0.540 | 22% |

**Fig. 5.**
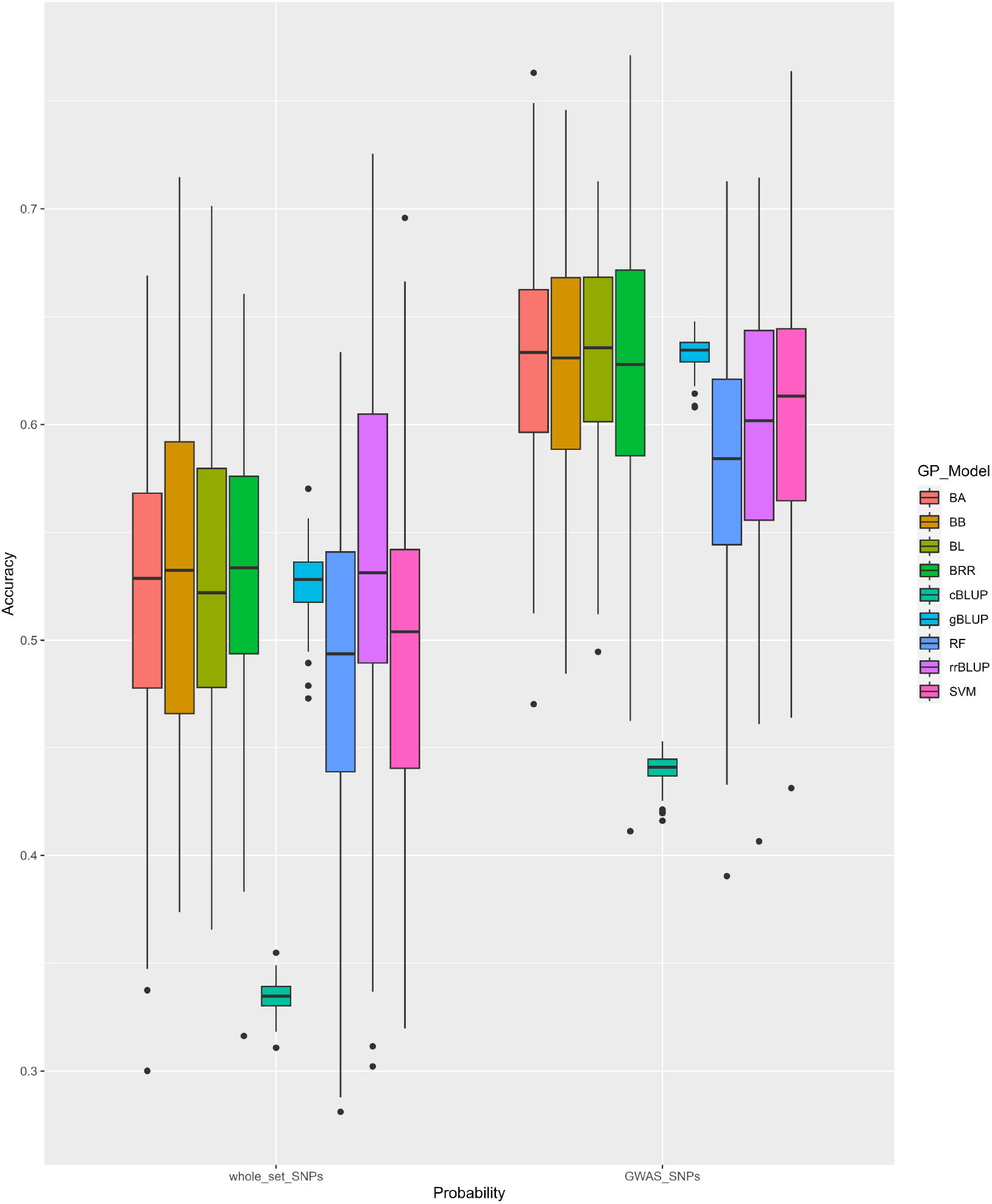
Genomic prediction for high sucrose contents using whole-set and GWAS-derived SNPs.

## Discussion

### Wider genetic diversity in soybean for improvement of seed sucrose concentration

Sucrose is desirable for the better taste and flavor of soy-derived foods. High sucrose concentration in soybean seeds is becoming an important breeding objective for development of food-grade soybean cultivars. To successfully meet the food-grade quality standards, seed sucrose concentration should be 6.0–8.0% (OSACC, 2020). During recent years, significant attention was given to characterising the soybean germplasm globally for exploring the range of available genetic diversity in seed sucrose concentration. For example, Sui et al. (2020) evaluated 178 genetically diverse soybean genotypes across three locations in China and reported 3.33–9.94% seed sucrose contents. Likewise, Ficht et al. (2022) characterized a panel of 266 historic and current soybean accessions across four field locations in Ontario (Canada) and described a wider genetic variation of 1.48–9.31% in seed sucrose concentration. Lu et al. (2022) assessed 278 diverse soybean accessions across three different locations and reported 5.52–16.46% total soluble sugar contents. Similarly, Xu et al. (2022) reported 0.58–13.99% total soluble sugars in 264 soybean accessions evaluated under two different environments. In this study also, we selected 473 USDA soybean germplasm accessions in which seed sucrose contents showed a near-symmetrical distribution with a wider genetic variation of 0.1–10.5% (**Fig. 1, Table S1**). Four Asian countries including Japan, South Korea, China and Russia harbored relatively diverse accessions, as large variations were observed in seed sucrose contents of soybean accessions originating from these countries. Among all accessions, at least 83 contained 6.0-10.5% sucrose concentration in their seeds, indicating these as “food-grade soybean” accessions (**Table S1**). Moreover, at least two accessions with >10% sucrose contents were also identified which could be exploited as donor parents for high sucrose contents in soybean breeding programs. Overall, the availability of this wider genetic diversity provides a rich resource for further improvement of seed sucrose concentration along with desirable agronomic and seed composition traits.

### Stable SNP markers for screening of soybean genotypes

Molecular markers are an essential prerequisite for accurate and robust screening of genetically diverse germplasm sets. Linkage analysis and GWAS are the most commonly used methods for molecular marker identification. To date, 42 QTLs have been reported for soybean seed sucrose contents through linkage analysis of bi-parental mapping populations (https://www.soybase.org/search/qtllist_by_symbol.php, accessed on February 13, 2023). However, only a few studies involving GWAS have been reported in recent years (Sui et al. 2020; Ficht et al. 2022; Xu et al. 2022; Lu et al. 2022). This study also followed GWAS approach to pinpoint important genomic regions significantly associated with seed sucrose contents. After subjecting the genetic and biochemical variation information of 8,477 high-quality SNPs and 473 USDA soybean germplasm accessions respectively, 75 SNPs were significantly (LOD ≥6.0) associated with seed sucrose contents (**Fig. 3A**). Among these, four stable SNP markers were also identified which showed overlapping between two or three GWAS tested models (**Table 1**). However, none of these stable SNPs overlapped with previously reported GWAS-derived SNPs indicating these as novel and stable for screening of diverse soybean germplasm sets. Possible reasons for their non-overlapping with already identified SNPs include a comparatively higher LOD threshold and differences in the number of high-quality SNP sets, genetic background of germplasm sets, GWAS-tested models and softwares/packages used for GWAS. Overall, on basis of a combination of highest LOD threshold and overlapping between GWAS-tested models, it could be expected that four candidate SNP markers identified in this study are comparatively more stable and efficient than previously reported GWAS-derived SNPs for screening of soybean germplasm accessions containing genetic variation in their seed sucrose contents.

### Candidate genes regulating sucrose biosynthesis, transport and storage

GWAS has become a mainstream approach for the identification of trait-linked genes due to involvement of relatively short linkage disequilibrium fragments (Li et al. 2015). At present, several hundreds of genes have been identified to participate in the regulation of soybean sucrose concentration (Sui et al. 2020; Lu et al. 2022). In this study, 23 candidate genes were identified within 20 kb flanking genomic regions of four stable SNPs (**Table 2**). Among these, at least 15 candidate genes were moderate to highly expressed in seeds and pod shells, along with other vegetative tissues (Severin et al. 2010). Interestingly, a sugar transporter encoding major facilitator superfamily gene (*Glyma*.*06G132500*) showing highest expression in pod shells at 10–17 DAF was also identified. Positioning of these candidate genes near four stable SNP peaks and their expression patterns in seeds and pods mimic their putative roles in regulating soybean seed sucrose concentration. Until now, none of these candidate genes has been functionally characterized. Their detailed characterization would be intriguing, however, beyond the scope of the current study, and could open new avenues to breed sucrose-enriched soybean. Therefore, these genes are worth exploring in future studies for deciphering their possible roles in sucrose biosynthesis, transport and storage within soybean plants.

### Marker-assisted development of sucrose enriched soybean cultivars

Marker-assisted breeding has proven greater advantages over conventional breeding for the improvement of plant quantitative traits. Among various molecular markers, kompetitive allele specific PCR (KASP) marker assays offer high-throughput accuracy, efficiency and low cost for the detection of SNPs in large and diverse germplasm sets (He et al. 2014). KASP markers have several applications in marker-assisted selection and breeding for the improvement of crop yield, resistance/tolerance and quality traits (Patil et al. 2016, 2017; Zhao et al. 2019; Liu et al. 2020; Wilkes et al. 2023). For example, Shi et al. (2021) designed a KASP marker assay which proved to be efficient for high-throughput selection of soybean cyst nematode-resistant lines. Similarly, Luciana Rosso et al. (2021) developed a breeder-friendly KASP assay linked to low Kunitz trypsin inhibitor (KTI) which showed 86% selection efficiency and could be used for marker-assisted breeding of low KTI concentration in soybean seeds. Likewise, Pan et al. (2022) developed a KASP marker of *Glyma*.*19G122500* and reported 90% and 66.33% selection efficiencies for screening of low and high soluble sugar containing soybean genotypes, respectively. In this study, selection accuracy and efficiency along with favorable alleles of GWAS-derived SNPs were estimated for marker-assisted selection and breeding of sucrose enriched soybean (**Table S4**). High selection accuracy (>66%) and efficiency (>40%) were computed for four stable SNPs, except for one SNP which showed comparatively moderate efficiency (24.5%). Moreover, their favorable alleles were also identified using contrasting soybean genotypes with high and low seed sucrose contents. However, development and validation of KASP markers assay are required, which is beyond the scope of this study, for practical utilization of identified SNP markers and their favorable alleles in soybean breeding programs. In future, additional studies involving development and validation of KASP markers could reveal their practicability in marker-assisted breeding of sucrose enriched soybean cultivars.

### Genomic selection/prediction for higher genetic gains

Genomic prediction is one of the most important parameters for measuring the performance of genomic selection. GP accuracy is dependent upon several factors including objective trait with its heritability, markers density and their association with the trait, population size, level of linkage disequilibrium, genomic selection models and relationship between training to testing populations (Shi et al. 2021). In this study, GP accuracy was tested using three scenarios: (1) different training to testing set, (2) different GP models, and (3) using whole-set versus GWAS-derived SNPs. The five training/testing sets (50/50, 67/33, 75/25, 83/17 and 90/10) showed similar but non-identical averaged r-values (range 0.34–0.52) (**Fig. 4**). Marginal improvements in GP accuracy (0.03–0.07) were observed when training/testing set ratios were increased (**Table S5**). The highest accuracy was observed in 90/10 ratio set, however, variance also enlarged. Moreover, GP accuracy testing with nine GP models also revealed similar results, although slightly lower accuracy values were observed with cBLUP model (**Fig. 4**). Comparable accuracy trends have been reported in other recent studies (Keller et al. 2020; Ravelombola et al. 2021; Shi et al. 2021, 2022). Similarly, testing with GWAS-derived SNPs showed significantly higher accuracy values across all GP models than whole-set SNPs (**Fig. 5**). On average, a 22% increase in prediction accuracy was observed which ranged from 16%– 33% (**Table 3**). GWAS-derived SNPs have also shown higher prediction accuracy values than whole-set and randomly selected SNPs for seed size and amino acid contents in soybean (Zhang et al. 2016; Qin et al. 2019). Collectively, our results highlighted that GWAS-derived SNPs are more efficient and any training/testing set ratio of ≥ 50% and tested GP models, except cBLUP, could be used for genomic selection/prediction of sucrose enriched soybean seeds. However, genomic predictions could be biased when GWAS-derived SNPs are used to predict genomics-assisted breeding values of the same GWAS panel or probably lower prediction accuracy may be observed in other panels with different individuals (Shi et al. 2022). In conclusion, genomic selection/prediction along with GWAS-derived SNPs and MAS would be beneficial for marker-assisted breeding of high sucrose-containing soybean cultivars and/ or other quantitative traits of important plant species.

## Conclusion

In this study, 473 USDA soybean germplasm accessions were evaluated for estimation of genetic diversity in seed sucrose contents. Wider genetic diversity existed in accessions originating from four Asian countries including China, Japan, Russia and South Korea. Tow accessions with >10% seed sucrose contents (PI398768 and PI229343) were identified which could be utilized as donor parents for the development of food-grade soybean cultivars. Population structure, PCA and phylogenetic analyses classified USDA germplasm accessions into three distinct groups or clusters having stable boundaries with well-defined genetic backgrounds. GWAS with 8,477 high-quality SNPs pinpointed four stable and large-effect SNPs/QTLs (Gm03_45385087_ss715586641, Gm06_10919443_ss715592728, Gm09_45335932_ss715604570 and Gm14_10470463_ss715617454) significantly associated (LOD ≥6.0) with seed sucrose contents. A total of 23 candidate genes including a sugar transporter (*Glyma*.*06G132500*) were identified from 20 kb flanking genomic regions of four stable SNPs. Selection accuracy and efficiency, as well as favorable alleles, of GWAS-derived SNPs were also estimated. Moreover, genomic predictions were tested using three different scenarios and results revealed that GWAS-derived SNPs are more valuable for the improvement of soybean seed sucrose contents through marker-assisted and genomic selections.

## Supplementary information

All supplementary information has been provided with this manuscript.

## Author contribution

AR and QR performed statistical analysis and wrote the draft of the manuscript. AK, DD, KC, TMP and AS assisted in data analysis. All authors have edited, reviewed, and approved the manuscript.

## Data availability

The datasets used and analyzed during the current study are available from the corresponding author upon reasonable request.

## References

Akond M, Liu S, Kantartzi SK, et al (2015) Quantitative Trait Loci Underlying Seed Sugars Content in “MD96-5722” by “Spencer” Recombinant Inbred Line Population of Soybean. Food Nutr Sci 6:964–973. https://doi.org/10.4236/FNS.2015.611100

Bellaloui N, Ebelhar MW, Gillen AM, et al (2011) Soybean seed protein, oil, and fatty acids are altered by S and S + N fertilizers under irrigated or non-irrigated environments. Agric Sci 2:465–476. https://doi.org/10.4236/AS.2011.24060

Bland JM, Altman DG (1995) Multiple significance tests: the Bonferroni method. BMJ 310:170. https://doi.org/10.1136/BMJ.310.6973.170

Breiman L (2001) Random forests. Mach Learn 45:5–32. https://doi.org/10.1023/A:1010933404324/METRICS

Cao Y, Li S, Wang Z, et al (2017) Identification of major quantitative trait loci for seed oil content in soybeans by combining linkage and genome-wide association mapping. Front Plant Sci 8:1222. https://doi.org/10.3389/FPLS.2017.01222/BIBTEX

Choung M-G (2010) Determination of Sucrose Content in Soybean Using Near-infrared Reflectance Spectroscopy. J Korean Soc Appl Biol Chem 53:478–484. https://doi.org/10.3839/jksabc.2010.073

Du Y, Zhao Q, Chen L, et al (2020) Effect of drought stress on sugar metabolism in leaves and roots of soybean seedlings. Plant Physiol Biochem 146:1–12. https://doi.org/10.1016/J.PLAPHY.2019.11.003

Endelman JB (2011) Ridge Regression and Other Kernels for Genomic Selection with R Package rrBLUP. Plant Genome 4:250–255. https://doi.org/10.3835/PLANTGENOME2011.08.0024

Fang C, Ma Y, Wu S, et al (2017) Genome-wide association studies dissect the genetic networks underlying agronomical traits in soybean. Genome Biol 18:1–14. https://doi.org/10.1186/S13059-017-1289-9/FIGURES/5

Farrar J, Pollock C, Gallagher J (2000) Sucrose and the integration of metabolism in vascular plants. Plant Sci 154:1–11. https://doi.org/10.1016/S0168-9452(99)00260-5

Ficht A, Bruce R, Torkamaneh D, et al (2022) Genetic analysis of sucrose concentration in soybean seeds using a historical soybean genomic panel. Theor Appl Genet 135:1375–1383. https://doi.org/10.1007/S00122-022-04040-Z/TABLES/5

Frichot E, François O (2015) LEA: An R package for landscape and ecological association studies. Methods Ecol Evol 6:925–929. https://doi.org/10.1111/2041-210X.12382

Grant D, Nelson RT, Cannon SB, Shoemaker RC (2010) SoyBase, the USDA-ARS soybean genetics and genomics database. Nucleic Acids Res 38:D843–D846. https://doi.org/10.1093/NAR/GKP798

He C, Holme J, Anthony J (2014) SNP genotyping: The KASP assay. Methods Mol Biol 1145:75–86. https://doi.org/10.1007/978-1-4939-0446-4_7/COVER

Heffner EL, Sorrells ME, Jannink JL (2009) Genomic Selection for Crop Improvement. Crop Sci 49:1–12. https://doi.org/10.2135/CROPSCI2008.08.0512

Hou A, Chen P, Alloatti J, et al (2009) Genetic Variability of Seed Sugar Content in Worldwide Soybean Germplasm Collections. Crop Sci 49:903–912. https://doi.org/10.2135/CROPSCI2008.05.0256

Huang M, Liu X, Zhou Y, et al (2019) BLINK: a package for the next level of genome-wide association studies with both individuals and markers in the millions. Gigascience 8:1–12. https://doi.org/10.1093/GIGASCIENCE/GIY154

Hwang EY, Song Q, Jia G, et al (2014) A genome-wide association study of seed protein and oil content in soybean. BMC Genomics 15:1–12. https://doi.org/10.1186/1471-2164-15-1/TABLES/3

Hymowitz T, Collins FI, Panczner J, Walker WM (1972) Relationship Between the Content of Oil, Protein, and Sugar in Soybean Seed1. Agron J 64:613–616. https://doi.org/10.2134/AGRONJ1972.00021962006400050019X

Karatzoglou A, Smola A, Hornik K (2023) kernlab: Kernel-Based Machine Learning Lab. R package version 0.9-32,https://CRAN.R-project.org/package=kernlab

Keller B, Ariza-Suarez D, de la Hoz J, et al (2020) Genomic Prediction of Agronomic Traits in Common Bean (Phaseolus vulgaris L.) Under Environmental Stress. Front Plant Sci 11:1001. https://doi.org/10.3389/FPLS.2020.01001/BIBTEX

Khan MA, Tong F, Wang W, et al (2019) Correction to: Analysis of QTL–allele system conferring drought tolerance at seedling stage in a nested association mapping population of soybean [Glycine max (L.) Merr.] using a novel GWAS procedure (Planta, (2018), 248, 4, (947-962), 10.1007/s00425-018-2952-4). Planta 249:1653. https://doi.org/10.1007/S00425-019-03143-0/METRICS

Kim HK, Kang ST, Cho JH, et al (2005) Quantitative trait loci associated with oligosaccharide and sucrose contents in soybean (Glycine max L.). J Plant Biol 48:106–112. https://doi.org/10.1007/BF03030569/METRICS

Kim HK, Kang ST, Oh KW (2006) Mapping of putative quantitative trait loci controlling the total oligosaccharide and sucrose content of Glycine max seeds. J Plant Res 119:533–538. https://doi.org/10.1007/S10265-006-0004-9/TABLES/4

Korte A, Farlow A (2013) The advantages and limitations of trait analysis with GWAS: A review. Plant Methods 9:1–9. https://doi.org/10.1186/1746-4811-9-29/FIGURES/4

Krober OA, Cartter -° JL (1962) Quantitative Interrelations of Protein and Nonprotein Constituents of Soybeans1. Crop Sci 2:171–172. https://doi.org/10.2135/CROPSCI1962.0011183X000200020028X

Lee C, Choi M-S, Kim H-T, et al (2015) Soybean [Glycine max (L.) Merrill]: Importance as A Crop and Pedigree Reconstruction of Korean Varieties. Plant Breed Biotechnol 3:179–196. https://doi.org/10.9787/PBB.2015.3.3.179

Lee T, Kim K Do, Kim JM, et al (2021) Genome□wide association study for ultraviolet□b resistance in soybean (Glycine max l.). Plants 10:1335. https://doi.org/10.3390/PLANTS10071335/S1

Li Y-S, Du M, Zhang Q-Y, et al (2012) Greater differences exist in seed protein, oil, total soluble sugar and sucrose content of vegetable soybean genotypes [Glycine max (L.) Merrill] in Northeast China. AJCS 6:1681–1686

Li Y hui, Reif JC, Ma Y song, et al (2015) Targeted association mapping demonstrating the complex molecular genetics of fatty acid formation in soybean. BMC Genomics 16:1–13. https://doi.org/10.1186/S12864-015-2049-4/FIGURES/5

Liu L, Song W, Wang L, et al (2020) Allele combinations of maturity genes E1-E4 affect adaptation of soybean to diverse geographic regions and farming systems in China. PLoS One 15:e0235397. https://doi.org/10.1371/JOURNAL.PONE.0235397

Liu X, Huang M, Fan B, et al (2016) Iterative Usage of Fixed and Random Effect Models for Powerful and Efficient Genome-Wide Association Studies. PLOS Genet 12:e1005767. https://doi.org/10.1371/JOURNAL.PGEN.1005767

Lu W, Sui M, Zhao X, et al (2022) Genome-Wide Identification of Candidate Genes Underlying Soluble Sugar Content in Vegetable Soybean (Glycine max L.) via Association and Expression Analysis. Front Plant Sci 13:1910. https://doi.org/10.3389/FPLS.2022.930639/BIBTEX

Luciana Rosso M, Shang C, Song Q, et al (2021) Development of Breeder-Friendly KASP Markers for Low Concentration of Kunitz Trypsin Inhibitor in Soybean Seeds. Int J Mol Sci 2021, Vol 22, Page 2675 22:2675. https://doi.org/10.3390/IJMS22052675

Lynch H, Johnston C, Wharton C (2018) Plant-Based Diets: Considerations for Environmental Impact, Protein Quality, and Exercise Performance. Nutr 2018, Vol 10, Page 1841 10:1841. https://doi.org/10.3390/NU10121841

Maroof MAS, Buss GR (2011) Low Phytic Acid, Low Stachyose, High Sucrose Soybean Lines. United States Patent Appl. Publ. U.S. Patent No. 8,003,856.

Maughan PJ, Maroof MAS, Buss GR (2000) Identification of quantitative trait loci controlling sucrose content in soybean (Glycine max). Mol Breed 6:105–111

Nelder JA, Wedderburn RWM (1972) Generalized Linear Models. J R Stat Soc Ser A 135:370–384. https://doi.org/10.2307/2344614

OSACC (2020) Ontario soybean and canola committee. Available online:http://www.gosoy.ca. Accessed 27 January 2023.

Pan W jing, Han X, Huang S yu, et al (2022) Identification of candidate genes related to soluble sugar contents in soybean seeds using multiple genetic analyses. J Integr Agric 21:1886–1902. https://doi.org/10.1016/S2095-3119(21)63653-5

Patil G, Chaudhary J, Vuong TD, et al (2017) Development of SNP Genotyping Assays for Seed Composition Traits in Soybean. Int J Plant Genomics 2017. https://doi.org/10.1155/2017/6572969

Patil G, Do T, Vuong TD, et al (2016) Genomic-assisted haplotype analysis and the development of high-throughput SNP markers for salinity tolerance in soybean. Sci Reports 2016 61 6:1–13. https://doi.org/10.1038/srep19199

Patil G, Vuong TD, Kale S, et al (2018) Dissecting genomic hotspots underlying seed protein, oil, and sucrose content in an interspecific mapping population of soybean using high-density linkage mapping. Plant Biotechnol J 16:1939–1953. https://doi.org/10.1111/PBI.12929

Pérez P, De Los Campos G (2014) Genome-wide regression and prediction with the BGLR statistical package. Genetics 198:483–495. https://doi.org/10.1534/GENETICS.114.164442/-/DC1

Poysa V, Woodrow L (2002) Stability of soybean seed composition and its effect on soymilk and tofu yield and quality. Food Res Int 35:337–345. https://doi.org/10.1016/S0963-9969(01)00125-9

Qin J, Shi A, Song Q, et al (2019) Genome Wide Association Study and Genomic Selection of Amino Acid Concentrations in Soybean Seeds. Front Plant Sci 10:1445. https://doi.org/10.3389/FPLS.2019.01445/BIBTEX

Qiu LJ, Chen PY, Liu ZX, et al (2011) The worldwide utilization of the Chinese soybean germplasm collection. Plant Genet Resour 9:109–122. https://doi.org/10.1017/S1479262110000493

Rao MSS, Mullinix BG, Rangappa M, et al (2002) Genotype × Environment Interactions and Yield Stability of Food-Grade Soybean Genotypes. Agron J 94:72–80. https://doi.org/10.2134/AGRONJ2002.7200

Ravelombola W, Shi A, Huynh BL (2021) Loci discovery, network-guided approach, and genomic prediction for drought tolerance index in a multi-parent advanced generation intercross (MAGIC) cowpea population. Hortic Res 2021 81 8:1–13. https://doi.org/10.1038/s41438-021-00462-w

Ravelombola WS, Qin J, Shi A, et al (2019) Genome-wide association study and genomic selection for soybean chlorophyll content associated with soybean cyst nematode tolerance. BMC Genomics 20:1–18. https://doi.org/10.1186/S12864-019-6275-Z/FIGURES/5

Ruan Y-L (2012) Signaling Role of Sucrose Metabolism in Development. Mol Plant 5:763–765. https://doi.org/10.1093/mp/sss046

Ruan Y-L, Jin Y, Yang Y-J, et al (2010) Sugar Input, Metabolism, and Signaling Mediated by Invertase: Roles in Development, Yield Potential, and Response to Drought and Heat. Mol Plant 3:942–955. https://doi.org/10.1093/mp/ssq044

R Core Team (2022) R: A language and environment for statistical computing. R Foundation for Statistical Computing, Vienna, Austria. URL https://www.R-project.org/

Schmutz J, Cannon SB, Schlueter J, et al (2010) Genome sequence of the palaeopolyploid soybean. Nat 2010 4637278 463:178–183. https://doi.org/10.1038/nature08670

Severin AJ, Woody JL, Bolon YT, et al (2010) RNA-Seq Atlas of Glycine max: A guide to the soybean transcriptome. BMC Plant Biol 10:1–16. https://doi.org/10.1186/1471-2229-10-160/TABLES/3

Shi A, Bhattarai G, Xiong H, et al (2022) Genome-wide association study and genomic prediction of white rust resistance in USDA GRIN spinach germplasm. Hortic Res 9. https://doi.org/10.1093/HR/UHAC069

Shi A, Buckley B, Mou B, et al (2016) Association analysis of cowpea bacterial blight resistance in USDA cowpea germplasm. Euphytica 208:143–155. https://doi.org/10.1007/S10681-015-1610-1/TABLES/2

Shi A, Gepts P, Song Q, et al (2021) Genome-Wide Association Study and Genomic Prediction for Soybean Cyst Nematode Resistance in USDA Common Bean (Phaseolus vulgaris) Core Collection. Front Plant Sci 12:1087. https://doi.org/10.3389/FPLS.2021.624156/BIBTEX

Skoneczka JA, Saghai Maroof MA, Shang C, Buss GR (2009) Identification of Candidate Gene Mutation Associated With Low Stachyose Phenotype in Soybean Line PI200508. Crop Sci 49:247–255. https://doi.org/10.2135/CROPSCI2008.07.0403

Song Q, Hyten DL, Jia G, et al (2015) Fingerprinting soybean germplasm and its utility in genomic research. G3: Genes genom genet 5(10), pp.1999–2006.

Sui M, Wang Y, Bao Y, et al (2020) Genome-wide association analysis of sucrose concentration in soybean (Glycine max L.) seed based on high-throughput sequencing. Plant Genome 13:e20059. https://doi.org/10.1002/TPG2.20059

Taira H, Tanaka H, Saito M, Saito M (1990) Effect of Cultivar, Seed Size, and Crop Year on Total and Free Sugar Contents of Domestic Soybeans. Nippon SHOKUHIN KOGYO GAKKAISHI 37:203–213. https://doi.org/10.3136/NSKKK1962.37.3_203

Teixeira AI, Ribeiro LF, Rezende ST, et al (2012) Development of a method to quantify sucrose in soybean grains. Food Chem 130:1134–1136. https://doi.org/10.1016/J.FOODCHEM.2011.07.128

Wang J, Zhang Z (2021) GAPIT Version 3: Boosting Power and Accuracy for Genomic Association and Prediction. Genomics Proteomics Bioinformatics 19:629–640. https://doi.org/10.1016/J.GPB.2021.08.005

Wang Y, Chen P, Zhang B (2014) Quantitative trait loci analysis of soluble sugar contents in soybean. Plant Breed 133:493–498. https://doi.org/10.1111/PBR.12178

Wickham H (2016) ggplot2: Elegant Graphics for Data Analysis. Springer-Verlag New York. ISBN 978-3-319-24277-4, https://ggplot2.tidyverse.org

Wilkes JE, Fallen B, Saski C, Agudelo P (2023) Development of SNP molecular markers associated with resistance to reniform nematode in soybean using KASP genotyping. Euphytica 219:1–10. https://doi.org/10.1007/S10681-022-03144-3/FIGURES/4

Wilson RF (2016) Seed Composition. Soybeans Improv Prod Uses 621–677. https://doi.org/10.2134/AGRONMONOGR16.3ED.C13

Xu W, Liu H, Li S, et al (2022) GWAS and Identification of Candidate Genes Associated with Seed Soluble Sugar Content in Vegetable Soybean. Agronomy 12:1470. https://doi.org/10.3390/AGRONOMY12061470/S1

Yang Y, Wang L, Zhang D, et al (2020) GWAS identifies two novel loci for photosynthetic traits related to phosphorus efficiency in soybean. Mol Breed 40:1–14. https://doi.org/10.1007/S11032-020-01112-0/METRICS

Yin L, Zhang H, Tang Z, et al (2021) rMVP: A Memory-efficient, Visualization-enhanced, and Parallel-accelerated Tool for Genome-wide Association Study. Genomics Proteomics Bioinformatics 19:619–628. https://doi.org/10.1016/J.GPB.2020.10.007

Zeng A, Chen P, Shi A, et al (2014) Identification of Quantitative Trait Loci for Sucrose Content in Soybean Seed. Crop Sci 54:554–564. https://doi.org/10.2135/CROPSCI2013.01.0036

Zhang J, Song Q, Cregan PB, Jiang GL (2016) Genome-wide association study, genomic prediction and marker-assisted selection for seed weight in soybean (Glycine max). Theor Appl Genet 129:117–130. https://doi.org/10.1007/S00122-015-2614-X/TABLES/3

Zhang Z, Ersoz E, Lai CQ, et al (2010) Mixed linear model approach adapted for genome-wide association studies. Nat Genet 2010 424 42:355–360. https://doi.org/10.1038/ng.546

Zhao J, Wang Z, Liu H, et al (2019) Global status of 47 major wheat loci controlling yield, quality, adaptation and stress resistance selected over the last century. BMC Plant Biol 19:1–14. https://doi.org/10.1186/S12870-018-1612-Y/FIGURES/6

Zhou Z, Jiang Y, Wang Z, et al (2015) Resequencing 302 wild and cultivated accessions identifies genes related to domestication and improvement in soybean. Nat Biotechnol 2015 334 33:408–414. https://doi.org/10.1038/nbt.3096

