## Supplementary figure for "GWAS and genomic selection for marker-assisted development of sucrose enriched soybean cultivars"

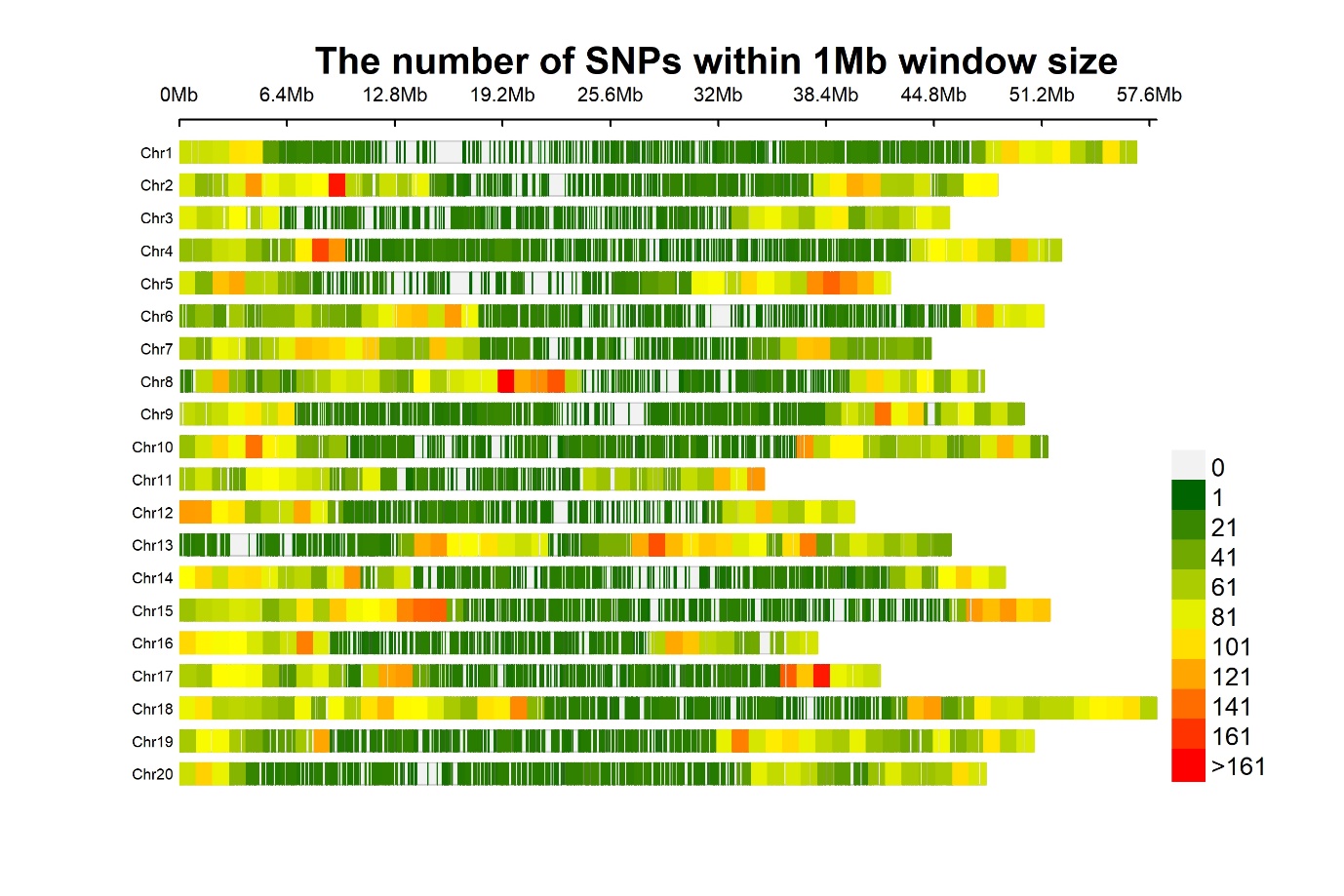

**Fig. S1** The density and distribution of single nucleotide polymorphism (SNP) across 20 soybean chromosomes

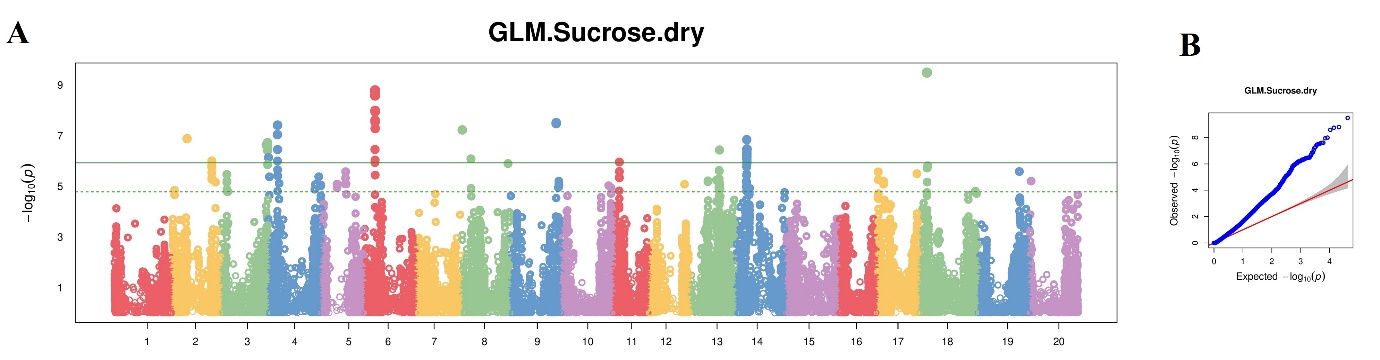

**Fig. S2** **(A)** Manhattan plot distribution of SNPs on 20 soybean chromosomes with GLM model. X- and y-axes indicate chromosomes and LOD values respectively. **(B)** QQ-plot distribution of observed vs expected LOD values

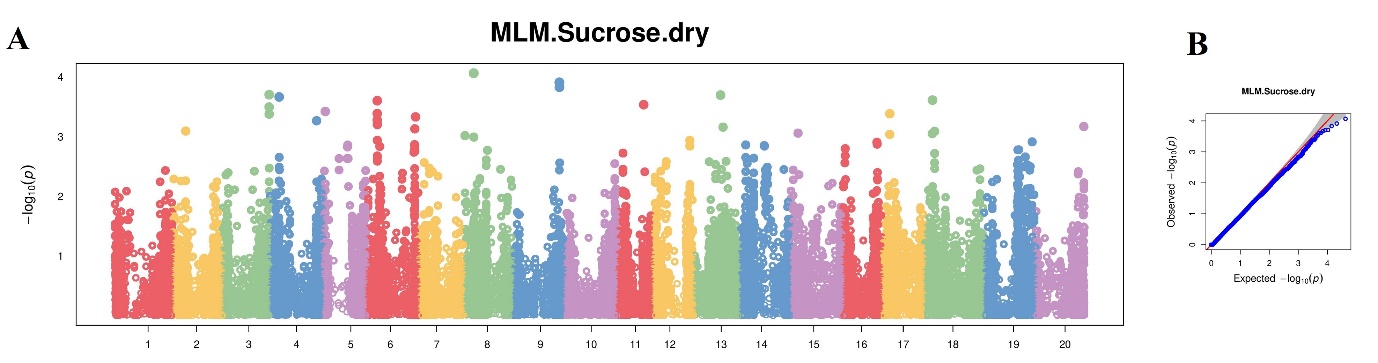

**Fig. S3** **(A)** Manhattan plot distribution of SNPs on 20 soybean chromosomes with MLM model. X- and y-axes indicate chromosomes and LOD values respectively. **(B)** QQ-plot distribution of observed vs expected LOD values

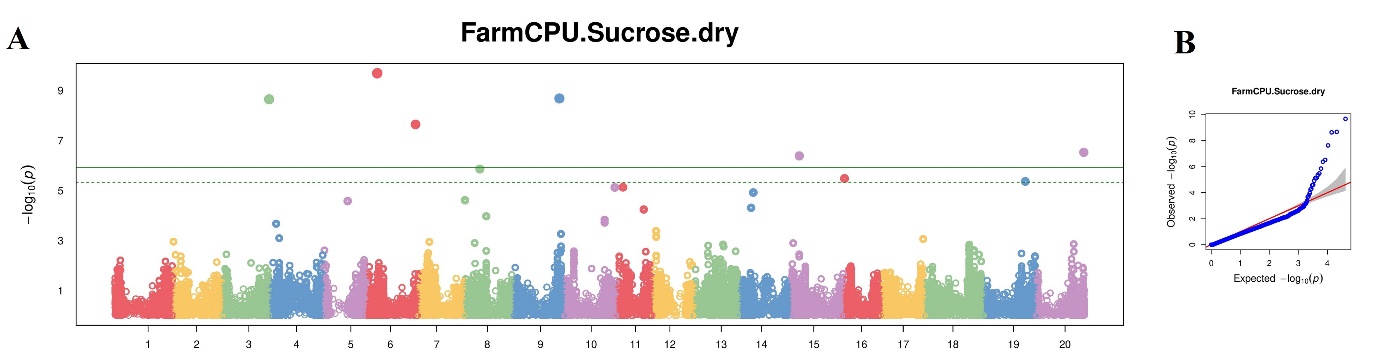

**Fig. S4** **(A)** Manhattan plot distribution of SNPs on 20 soybean chromosomes with FarmCPU model. X- and y-axes indicate chromosomes and LOD values respectively. **(B)** QQ-plot distribution of observed vs expected LOD values

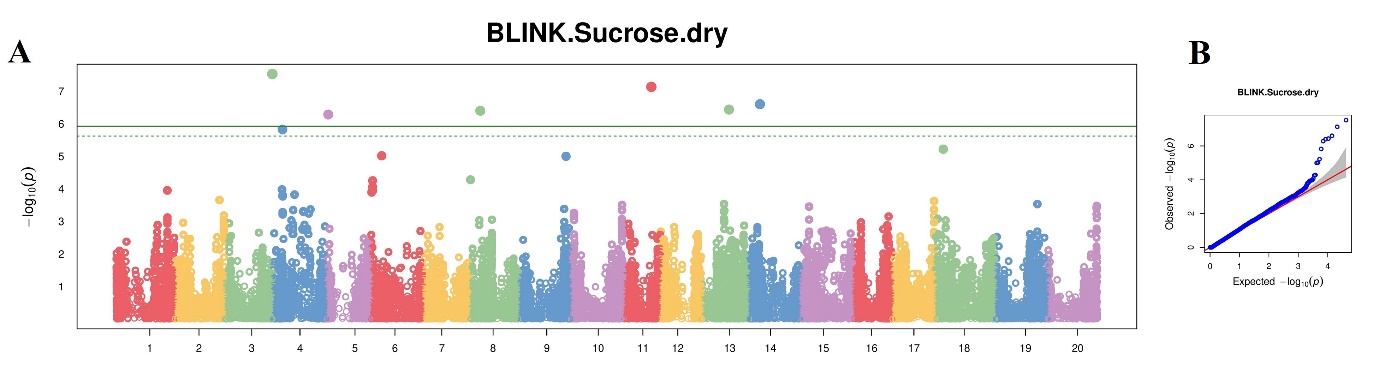

**Fig. S5** **(A)** Manhattan plot distribution of SNPs on 20 soybean chromosomes with BLINK model. X- and y-axes indicate chromosomes and LOD values respectively. **(B)** QQ-plot distribution of observed vs expected LOD values
